# Chromosome-level genome assembly of the gerbera (*Gerbera hybrida*) using HiFi long-read and Hi-C technologies

**DOI:** 10.1101/2025.08.15.670457

**Authors:** Yuta B. Aoyagi, Riki Shimada, Hideki Hirakawa, Atsushi Toyoda, Hidehiro Toh, Sachiko Isobe, Naoyuki Tajima, Kenta Shirasawa, Tokumasa Horiike, Hiroshi Fujii, Tao Fujiwara, Masaru Bamba, Takashi Nakatsuka, Akiyoshi Tominaga

## Abstract

*Gerbera hybrida* is one of the most popular ornamental plants and also serves as a valuable model plant within the Asteraceae family. Here, we report both the nuclear and organellar genome assemblies and annotations of *G. hybrida*, which was developed through hybridization of two wild species. Sequencing was performed using a combination of PacBio high fidelity (HiFi) reads and chromatin capture reads (Omni-C). The total span of the nuclear genome assembly is 2.32 gigabases, and 99.3% of the sequence assembled into 25 scaffolds, consistent with the known chromosome number. Genome annotation of the nuclear genome identified 36,160 protein-coding genes and 11,572 non-coding transcripts. The mitochondrial genome had 363,511 bp and contains 36 protein-coding genes, 3 rRNAs, and 21 tRNAs, while the chloroplast genome is 151,898 bp in length and includes 85 protein-coding genes, 8 rRNAs, and 37 tRNAs. This reference genome provides a foundational resource for future molecular breeding and genetic research in *Gerbera* and the broader Asteraceae family.

## 1. Introduction

*Gerbera hybrida* (2*n* = 2*x* = 50) displays an extensive range of flower colors including white, yellow, orange, peach, red, and purple, as well as various floral morphologies such as double, pasta type with twisted or curled petals, and spider types with long, narrow petals. It is among the most widely cultivated ornamental plants globally, with its distinctive floral structure and vibrant colors attracting many admirers.^1^ In Japan, the total shipment of gerbera cut flowers in 2023 reached 119.9 million units, ranking fourth after chrysanthemums, carnations, and roses.^2^ Most currently available commercial gerbera cultivars have been developed through interspecific hybridization, primarily in the Netherlands.^3^ *G. hybrida* is known to have originated from hybridization between two wild African species, *G. jamesonii* and *G. viridifolia*, although there are no detailed records of the hybridization process.^4^

The Asteraceae, to which genus *Gerbera* belongs, represents the largest family of flowering plants, comprising an estimated 25,000 to 35,000 species and accounting for approximately 10% of all angiosperms.^5^ The cephalic inflorescences characteristic of all Asteraceae are a fascinating subject of study, as they are a key factor contributing to the global success of this family. A defining feature of this family is the capitulum-type inflorescence, a key innovation contributing to the ecological and evolutionary success of the group.^6^ Another characteristic trait of the Asteraceae is the pappus, a highly modified calyx that facilitates seed dispersal and provides protection against herbivory by small animals.^7, 8^ Recent advancements in sequencing technologies have enabled the publication of whole-genome sequences for various Asteraceae species, including chrysanthemum,^9^ sunflower,^10^ lettuce,^11^ and *Mikania micrantha*, commonly known as the “mile-a-minute” weed.^12^ Furthermore, the chromosome-level genome assembly of *Chrysanthemum seticuspe* has been completed and is now widely used as a model for cultivated chrysanthemum research.^13^ *Gerbera* belongs to the subfamily Mutisioideae, which diverged early within the Asteraceae, and is phylogenetically distant from other Asteraceae species whose nucleotide sequences have been determined to date.^14, 15^ To date, no assembled genome has been reported for any member of the Mutisioideae. Therefore, whole-genome information of *Gerbera* is expected to provide valuable insights into the early evolutionary history of the Asteraceae.

*G. hybrida* serves as an important ornamental crop with significant global market demand. Its diploid genome makes it a useful model for investigating traits characteristic of the Asteraceae, such as capitulum inflorescence structure and metabolic diversity.^16, 17^ Several aspects of gerbera biology have already been examined, including petal development,^18, 19^ anthocyanin biosynthesis,^20, 21^ and male sterility.^22^ Additionally, unique secondary metabolite biosynthetic pathways in gerbera are important for understanding the evolution of environmental adaptation.^23^ Nonetheless, genomic resources for gerbera remain limited.

Currently, no publicly available genome data exist, and the linkage map constructed using single nucleotide polymorphism (SNP) markers comprises 27 linkage groups, two more than the haploid chromosome number.^24^ Recently, the brassinazole-resistant transcription factor family was identified based on whole-genome sequence data; however, these genome data have not yet been made publicly accessible.^25^ A comprehensive whole-genome assembly is urgently needed to support further molecular and evolutionary research in gerbera.

In this study, we generated a chromosome-level genome assembly of *G. hybrida* using long-read sequencing combined with Hi-C technologies. These genomic resources will facilitate molecular breeding of gerbera for traits such as flower morphology, pigmentation, and disease resistance, and will also provide a foundation for understanding the molecular mechanisms and evolutionary processes underlying key characteristics of Asteraceae.

## 2. Materials and methods

### 2.1 Library construction and sequencing

Fresh leaves and flower buds of mature *G. hybrida* ‘Opal’ were collected from temperature-controlled greenhouses at the Center for Education and Research in Field Science, Faculty of Agriculture, Shizuoka University (Figure 1). Samples were immediately frozen in liquid nitrogen and stored at –80 ℃ for further analysis.

**Figure 1.**
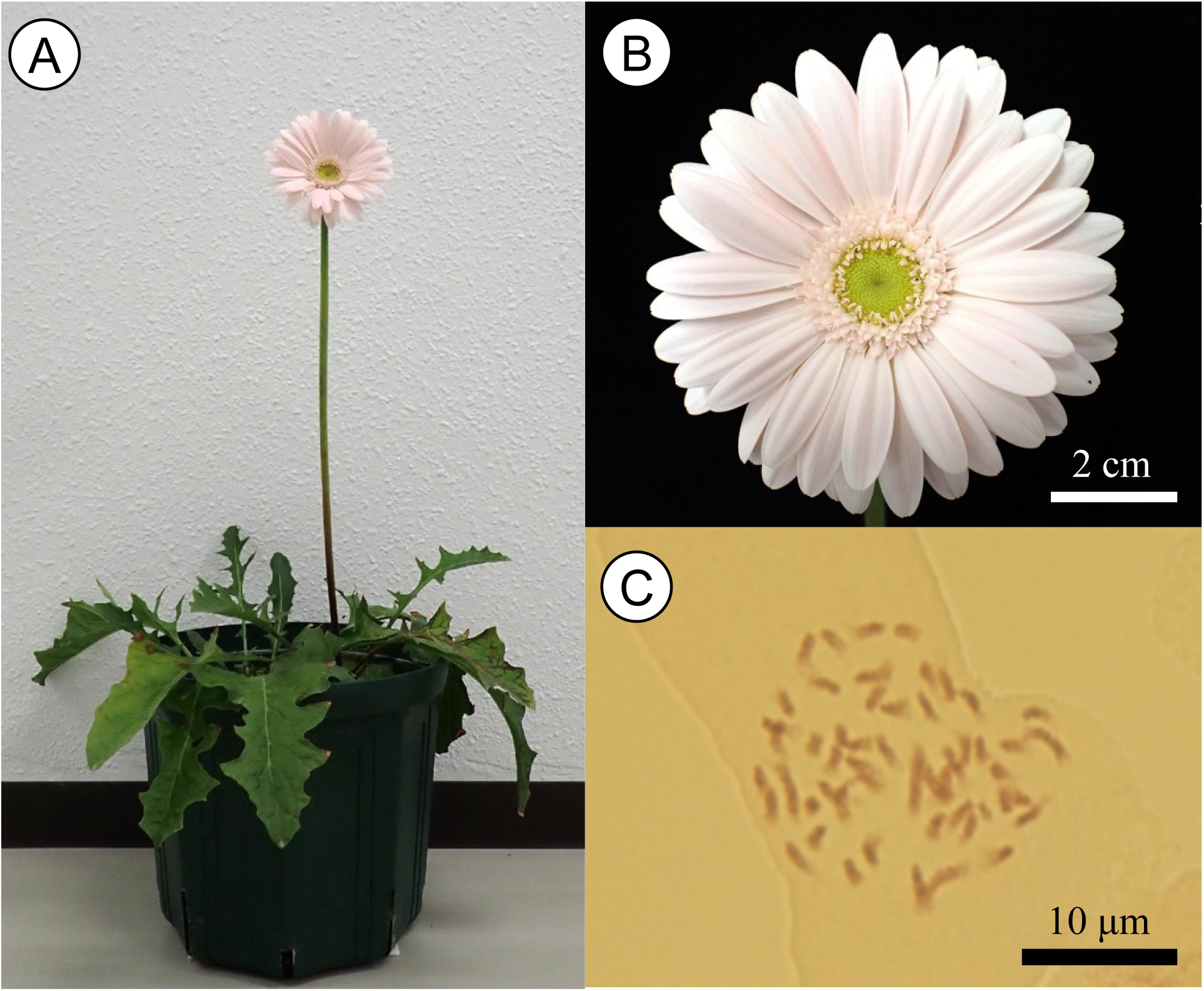
Morphology of *G. hybrida*. (A) Whole plant in flower, (B) Capitulum inflorescence, (C) metaphasic chromosome number 2*n*=2*x*=50.

Genomic DNA was extracted from leaves using the CTAB method and further purified with the Genomic-tip (QIAGEN). For long-read sequencing, the genomic DNA was sheared using a g-TUBE (Covaris) and a library was constructed using the SMRTbell prep kit 3.0 (Pacific Biosciences). DNA fragments of 15–20 kilobases (kb) were removed using the Sage ELF (Sage Science). Genome sequencing was performed on the Sequel IIe (Pacific Biosciences) using the Sequel II Binding Kit 3.2 and the Sequel II Sequencing Kit 2.0 (Pacific Biosciences). Six SMRT Cell 8M (Pacific Biosciences) were used, with a runtime of 30 hours per cell. HiFi reads were generated using the DeepConsensus v1.2.^26^ To generate Omni-C short reads, library was constructed using the Omni-C Kit (Dovetail). The library was sequenced on the NovaSeq 6000 (Illumina) using a 150 paired-end sequencing strategy.

Total RNA was extracted from leaves and flower buds using RNeasy Mini Kit (Qiagen). The extracted RNA was subsequently treated with DNase I (New England Biolabs) to remove genomic DNA contamination. cDNA libraries were constructed using the Illumina TruSeq Stranded mRNA HT Sample Prep Kit (Illumina). Sequence analysis was performed using the DNBSEQ-G400RS (MGI Tech) following the TruSeq Stranded mRNA Sample Preparation Guide.

### 2.2 Chromosome counts

Fresh thick roots were collected and pretreated in 2 mM 8-hydroxyquinoline solution at 15° C for 8 hours. They were then transferred to fixative solution (glacial acetic acid: ethanol = 1:3, v/v) at 4°C for more than 30 minutes. Next, the samples were placed in a mixture of 1% aceto-orcein solution and 1 N hydrochloric acid (9:1) and left to stand for 4 hours. Subsequently, they were treated with a mix of 9 drops of 2% aceto-orcein and 1 drop of 1 N HCl for 20 minutes. Finally, the squash method was applied using aceto-orcein solution as the staining reagent for microscopic observation.

### 2.3 Sequence read data processing

For HiFi reads, HiFi reads with remnant PacBio adapter sequences were removed using HiFiAdapterFilt v2.0.1^27^ with default parameters. For Omni-C reads, the bridge adapter sequences were trimmed from paired-end Omni-C reads using Cutadapt v3.5.^28^ Then, Illumina adapter sequences were trimmed, and reads with a base quality score less than 15 and a length shorter than 30 bases were filtered using fastp v0.12.4.^29^ For RNA sequencing (RNA-seq) reads, Illumina adapter sequences were trimmed, and reads with a base quality score less than 15 and a length shorter than 50 bases were filtered using fastp.

### 2.4 Genome size estimation

The haploid genome size of *G. hybrida* was estimated by counting the *k*-mer frequency of HiFi reads. 21-mers were counted using Meryl v1.3^30^ with default parameters. The genome size was then estimated using GenomeScope 2.0^31^ with ‘-p 2’ for the diploid genome, which was a parameter for ploidy in the model.

### 2.5 De novo genome assembly and scaffolding using Omni-C

HiFi reads were assembled with paired-end Omni-C reads using Hifiasm v0.19.5.^32, 33^ We used the primary assembly of Hifiasm for the *G. hybrida* genome assembly. Haplotigs were removed from the assembled genome sequences. HiFi reads were aligned to the assembled sequences using minimap2 v2.24-r1122^34, 35^ with ‘-x map-hifi’ and ‘--secondary=no’. Then, haplotigs were removed using Purge Haplotigs v1.1.2.^36^

After the purging haplotigs, scaffolding was performed using Omni-C. Alignments of Omni-C reads were generated following the Dovetail Omni-C’s mapping pipeline (https://omni-c.readthedocs.io/en/latest/). Omni-C reads were aligned to the assembled contigs using BWA v0.7.17-r1188^37^ with ‘mem-5-S-P-T0’. Alignments with a mapping quality of less than 40 were filtered, and PCR duplicates were removed using Pairtools v1.0.2.^38^ Then, the alignment result of Omni-C reads was used to correct for misjoin and orient the contigs into scaffolds using SALSA v2.3^39, 40^ with default parameters.

Subsequently, the scaffolds of organelle genomes were removed from the genome assembly. HiFi reads were assembled to generate the chloroplast and the mitochondrial genomes (see ‘Organelle genome assembly and annotation’ below for details). The scaffolds of the genome assembly were mapped to the chloroplast and the mitochondrial genome assemblies using minimap2. For each scaffold, alignment length was calculated using BEDTools.^41^ Then, the alignment coverage, where the sum of alignment lengths was divided by the contig length, was calculated. Finally, scaffolds with the alignment coverage greater than 0.95 were removed.

To correct scaffolding errors and remove redundant sequences, we manually corrected the scaffolds based on four types of information; (1) a contact map of Omni-C, (2) the HiFi read alignment, (3) syntenies between haploid assemblies generated by Hifiasm, and (4) the finding of telomeric repeats. The contact map of Omni-C was generated using Juicer v1.6.^42^ HiFi reads were aligned using minimap2 with ‘-x map-hifi’, and the alignment depth was calculated using deepTools bamCoverage.^43^ The scaffolds of the primary assembly were aligned to two haploid assemblies generated by Hifiasm using minimap2 with ‘-x asm5’, and a dot plot was created using D-Genies v1.4.0.^44^ Additionally, the plant-type telomeric repeat (TTTAGGG) was explored using a Telomere Identification toolKit (tidk) v0.2.31^45^ with default parameters. We cut misjoined scaffolds, where two contigs were joined between telomeric regions, based on the finding of telomeric repeats. We also removed redundant sequences based on the Omni-C contact map, the alignment depth of HiFi reads, and syntenic similarity between haploid assemblies. Then, based on the contact map and the syntenic similarity, we manually joined the scaffolds with 10,000 Ns. After curation, the genome assembled sequences were ordered by length. For the final genome assembly of *G. hybrida*, an Omni-C contact map was regenerated based on the alignment of Omni-C reads to the final scaffold sequences, and the telomeric repeat was explored by the method described above.

To assess the quality of the final genome assembly of *G. hybrida*, we quantified the completeness of the benchmarking universal single-copy orthologues (BUSCO) using BUSCO v5.7.1^46^ and the Embryophyta gene set from OrthoDB v10. In addition, to evaluate the assembly continuity based on the identification of long terminal repeat (LTR) retrotransposons, we calculated the LTR assembly index (LAI)^47^ for the assembled genome sequence using LTR_retriever v2.9.5.^48^

### 2.6 Repetitive sequence analysis

Repetitive sequences in the assembled genome sequences of *G. hybrida* were detected using RepeatModeler2 v2.0.5^49^ with ‘-LTRStruct’, which run the additional LTR search pipeline, and ‘-genomeSampleSizeMax=1G’. The custom repetitive sequence library was created by combining repetitive sequences detected by RepeatModeler and known repetitive sequences from Repbase^50^ and Dfam v3.8^51^ using RepeatMasker v4.1.5 (http://repeatmasker.org). Repetitive sequences in the assembled genome sequence were then annotated using RepeatMasker with the custom library and the ‘-s’ parameter, which enables a more sensitive search.

For the gene prediction described below, repetitive sequences in the assembled genome sequence were soft-masked with the library excluding repetitive sequences that were part of repetitive protein families by performing a BLASTX search against the UniProtKB database^52^ (accessed on July 3, 2023) using DIAMOND v2.1.8.162.^53^

### 2.7 Gene prediction and functional annotation

The protein-coding genes prediction was conducted for the repeat-masked assembled genome sequences by combining ab initio prediction and homology-based prediction. Ab initio prediction was performed using BRAKER3 v3.0.7^54–56^ with the alignment of RNA-seq reads by STAR v2.7.10b^57^ 2-pass mode and protein sequences in Viridiplantae from OrthoDB v12.^58^ In the BRAKER3 prediction, we set ‘--gc_probability’, a parameter of the probability for the donor splice site pattern GC in the gene prediction, to 0.01 based on the RNA-seq alignment information. For the homology-based prediction, we collected protein sequences of three published Asteraceae genomes listed in Table S1; *Mikania micrantha* (ASM936387v1),^12^ *Helianthus annuus* (HanXRQr2.0-SUNRISE),^10^ and *Lactuca sativa* (Lsat_Salinas_v11).^11^ The homology-based prediction was performed using GeMoMa v1.9^59, 60^ with three Asteraceae protein sequences and the alignment of RNA-seq reads from *G. hybrida*. For both BRAKER3 and GeMoMa predictions, for genes with multiple alternative transcripts, the longest transcript was determined as the representative transcript.

The predicted genes are considered to contain false positives. To generate a high-confidence gene set, we classified the predicted genes based on (1) the homology evidence and (2) the gene expression evidence, for BRAKER3 and GeMoMa predictions, respectively. For the homology evidence, BLASTP search of predicted genes was performed against the UniProtKB database^61^ with *E*-value less than 1e–20 and identity greater than 25% using DIAMOND in the more sensitive mode. The predicted genes were also searched against EggNOG v5.0 database^62^ with *E*-value less than 1e–10 using EggNOG-mapper v2.1.8.^63^ Domains in predicted genes were searched against Pfam v37.4,^64^ the protein domain database, with *E*-value less than 1e–10 using HMMER v3.3.2.^65^ For the gene expression evidence, RNA-seq reads were mapped to the predicted genes and the expression values in transcripts per million (TPM) were calculated using Salmon v1.10.2.^66^ Then, we classified the predicted genes into three categories; high-confidence (HC), low-confidence (LC), and transposable element-related (TE) genes. The genes with TPM value higher than 1 and those that had hits against databases were classified into HC genes. The genes that had hits against databases with the keywords related to TEs were classified into TE genes. The remaining genes were classified as LC genes. To create highly complete gene sets, we subsequently integrated HC genes from BRAKER3 and GeMoMa predictions. HC genes from BRAKER3 prediction were used as the reference, and structures of HC genes from GeMoMa and those from BRAKER3 were compared using GffCompare v0.12.6.^67^ For genes with a complete and/or partial match between the reference and the query, the reference genes and its structures were determined as representative genes. Furthermore, query genes that did not overlap with reference genes, those that were fully contained within a reference intron, and those that contained a reference gene within its intron were added to the representative genes. Finally, the quality and completeness of the predicted gene set was evaluated using BUSCO in protein mode.

In addition to protein-coding genes, we performed the prediction for non-coding RNA (ncRNA) genes, including tRNA, rRNA, microRNA (miRNA), small nuclear RNA (snRNA), and small nucleolar RNA (snoRNA). tRNAs were predicted using tRNAscan-SE 2.0 v2.0.12.^68^ We then conducted ‘EukHighConfidenceFilter’ in tRNAscan-SE 2.0 to filter out possible pseudogenes and tRNA genes with low-fidelity. Other types of RNA genes were searched using covariance models by Infernal v1.1.5^69^ with Rfam 15.0, a database of ncRNA families derived from a wide range of species.^70^ We selected 889 accessions from Rfam 15.0 that matched Viridiplantae sequences. RNA genes with an *E-*value less than 1e–5 were filtered. If different sequences from Rfam hit in the single genome region, we retained the sequence with the lowest *E-*value, or if multiple sequences had the same *E-*value, we retained the one with the highest bit score.

Circular plot of the *G. hybrida* genome assembly was visualized using Circos v0.69-8.^71^

### 2.8 Organelle genome assembly and annotation

To construct the chloroplast and the mitochondrial genomes of *G. hybrida*, HiFi reads were assembled using an organelle genome assembly toolkit (Oatk) v1.0^72^ with “-c 300”, which was the syncmer coverage threshold. Then, protein-coding, rRNA, and tRNA genes in the chloroplast and the mitochondrial genomes were annotated through GeSeq v2.0.3^73^ with a default parameter, respectively. For the chloroplast genome, protein-coding and rRNA genes were annotated based on BLAT^74^ with the MPI-MP chloroplast reference set, a manually curated reference set of chloroplast sequences provided by GeSeq. tRNA genes were annotated based on de novo prediction using ARAGORN.^75^ For the mitochondrial genome, protein-coding and rRNA genes were annotated based on BLAT with the mitochondrial sequences from eight Asteraceae species in NCBI RefSeq (Table S2). tRNA genes were annotated using ARAGORN.^75^ Finally, the structure of each gene was manually corrected based on the reference annotations. Organelle genome annotations were visualized by OGDRAW.^76^

Alignment and variant detection of chloroplast genome sequences were performed using the MUMmer4 package (v4.0.0beta2).^77^ The nucmer program was used to align *G. jamesonii* and *G. piloselloides* to the *G. hybrida* reference genome. The resulting alignments were filtered to retain the best one-to-one matches using “delta-filter-1”, and SNPs and InDels were extracted using “show-snps-Clr”.

### 2.9 Phylogenetic analysis and genome synteny

We collected protein sequences from publicly databases for *M. micrantha*, *H. annuus*, and *L. sativa*, as well as *Chrysanthemum seticuspe* (CSE_r1.0),^78^ *Artemisia annua* (ASM311234v1),^79^ *Taraxacum* kok-saghyz (OSU_TarKok_2.0), *Arctium lappa* (ASM2352574v1),^80^ and *Arabidopsis thaliana* (TAIR10.1).^81^ We constructed a species tree using a concatenated phylogenetic analysis based on a dataset of orthologous protein sequence identified by OrthoPhy.^82^ Subfamily-level taxonomic information was provided to OrthoPhy, classifying the nine Asteraceae species into four subfamilies: Asteroideae, Cichorioideae, Carduoideae, and Mutisioideae. An ortholog group contained at least one species in each subfamily. The tree was inferred using the maximum likelihood method with the LG substitution model.

Synteny analysis was performed using the MCScanX^83^ pipeline as follows. An all-versus-all protein BLASTP^84^ was conducted with an *E*-value cutoff of 1e–5, self-matches were removed, and syntenic blocks were detected with MCScanX using a minimum of 15 collinear gene pairs and allowing at most one intervening gene between collinear pairs. Finally, syntenic blocks were visualized using AccuSyn.^85^

## 3. Results and discussion

### 3.1 Nuclear genome assembly

In total, we obtained 254.34 gigabases (Gb) of PacBio HiFi reads with an N50 length of 18 kb, and 256.59 Gb of Dovetail Omni-C paired-end reads for *G. hybrida* (Table S3). In addition, the mRNA was obtained from 12 samples of *G. hybrida*, and the total length of paired-end RNA-seq reads was 41.18 Gb (Table S3).

HiFi reads were used for the estimation of the genome size and de novo genome assembly of *G. hybrida*. Based on the *k*-mer analysis, the haploid genome size was estimated to be 2.40 Gb, and the heterozygosity rate was estimated to be 2.05% (Figure S1). The C-value of *G. hybrida* by flow cytometry is 2.56 pg.^86^ This corresponds to about 2.36 Gb, very close to the estimate by *k*-mer analysis. It also suggests that *G. hybrida* has a complex genome with high heterozygosity, supporting that it is an interspecific hybrid between *G. viridifolia* and *G. jamesonii*.^4^

A total of 254.34 Gb (∼107× coverage) of HiFi reads and Omni-C reads were assembled, generating 2,138 primary contigs with a total length of 2,599.54 megabases (Mb) and an N50 of 83.20 Mb (Table S4). The BUSCO completeness of the primary assembly was 98.7% (89.7% of single-copy and 9.0% of duplicated BUSCOs). The relatively large assembly size compared to the estimated genome size and the high duplicated BUSCO score suggested the presence of remaining haplotigs in the genome assembly. We then removed haplotigs from the genome assembly. A total of 1,629 contigs were removed as haplotigs or artifactual contigs. After purging haplotigs, the genome assembly comprised 509 contigs with a total length of 2,498.20 Mb and 98.7% of BUSCO completeness (89.8% of single-copy and 8.9% of duplicated BUSCOs). The score of duplicated BUSCOs did not decrease significantly, indicating the incomplete removal of haplotigs.

After purging haplotigs, we conducted the scaffolding using Omni-C, elimination of organelle genome contigs, and the manual curation to remove remaining haplotigs in the genome assembly. These improved the continuity and reduced the redundancy of the genome assembly, resulting in the final *G. hybrida* genome assembly with a total length of 2,317.11 Mb and an N50 of 87.58 Mb (Figure 2A and Table 1). There were 25 scaffolds longer than 10 Mb, which corresponds to the number of chromosomes (Figure 1C and Table S5). These 25 scaffolds represent 99.3% of the entire assembly, and the fact that the interactions are found on the same chromosome in the contact map indicates a high quality of genome assembly (Figure 2B). Telomeric repeat (TTTAGGG) peaks were found at both ends of 19 scaffolds and at the single end of six scaffolds of the 25 scaffolds (Figure 2A). The BUSCO completeness of the final genome assembly was 98.7% (91.9% of single-copy and 6.8% of duplicated BUSCOs), which remained high completeness and decreased duplicated score after the manual curation. The whole genome LAI, a metric of the genome assembly continuity based on LTR retrotransposons, in the final genome assembly was 24.44, corresponding to gold quality (Table 1 and Figure S2).

**Figure 2.**
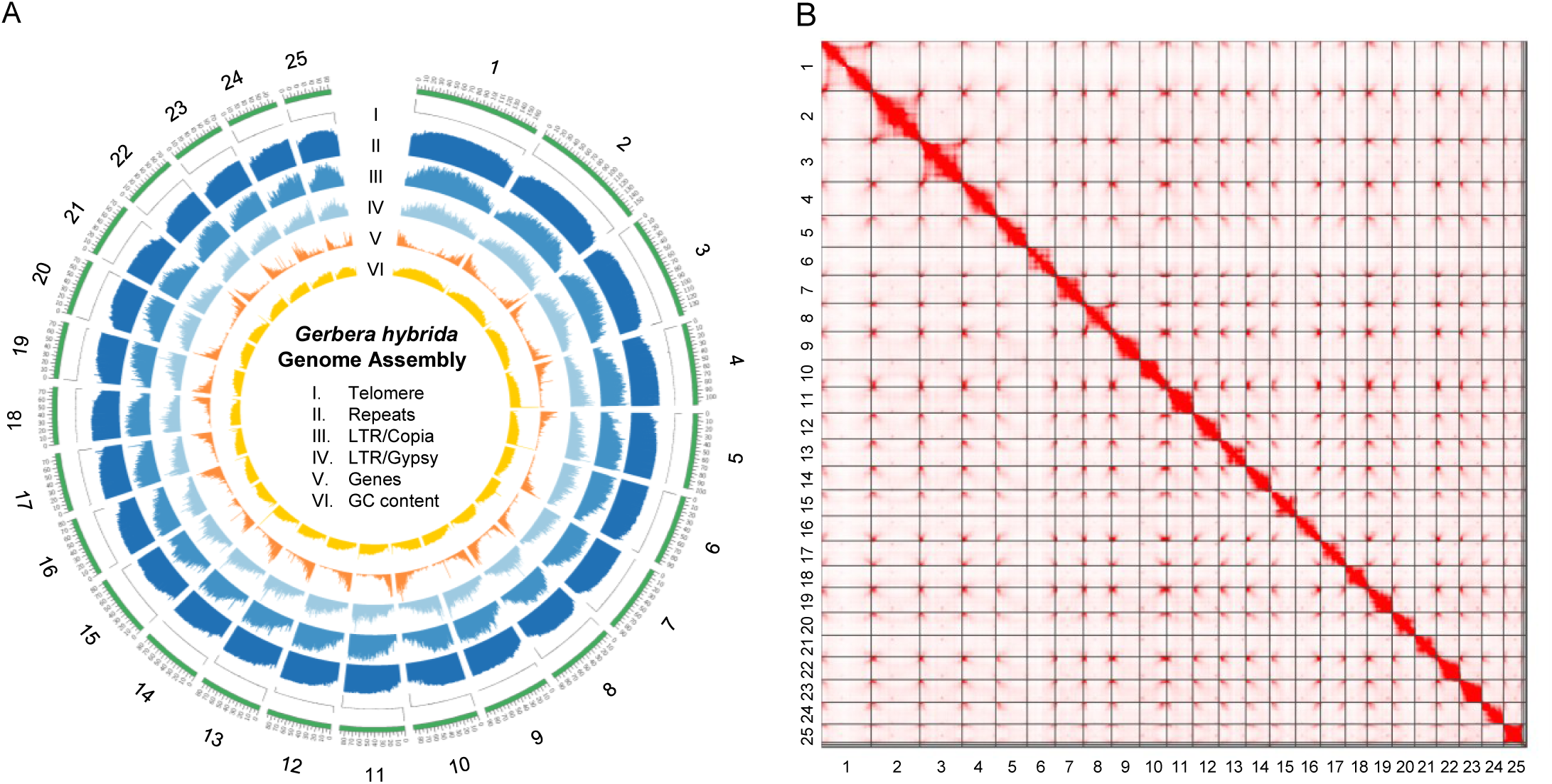
The genome assembly of *G. hybrida*. (A) The landscape of the *G. hybrida* genome assembly. Twenty-five sequences are shown. The track represents from outside to inside: (I) The count of identified telomeric repeats in a 10 kb window, with positive values representing the count of repeats in the reverse strand and negative values representing those in the forward strand, (II) The length of all repetitive sequences in a 500 kb window, (III) The length of Copia-type of LTR-retrotransposons in a 500 kb window, (IV) The length of Gypsy-type of LTR-retrotransposons in a 500 kb window, (V) The number of protein-coding genes in a 500 kb window, and (VI) The GC content in a 500 kb window. (B) The Omni-C contact map of the *G. hybrida* genome assembly. The numbers on the axes represent the scaffold number. A darker red color indicates higher interaction frequency.

**Table 1.** Statistics of the genome assembly of *G. hybrida*.

|  |  |
| --- | --- |
| Number of sequences | 42 |
| Total length (bases) | 2,317,114,601 |
| Average length (bases) | 55,169,395 |
| Maximum length (bases) | 169,236,808 |
| Minimum length (bases) | 23,246 |
| N50 (bases) | 87,579,574 |
| L50 | 11 |
| N90 (bases) | 72,510,611 |
| L90 | 22 |
| Number of gaps | 4 |
| Gap length (bases) | 21,000 |
| A (bases) | 727,623,957 |
| T (bases) | 728,075,646 |
| G (bases) | 430,627,287 |
| C (bases) | 430,766,711 |
| GC content (%) | 37.2 |
| BUSCO<sup>a</sup> |  |
| Complete (%) | 98.7 |
| (Complete and single-copy [%]) | 91.9 |
| (Complete and duplicated [%]) | 6.8 |
| Fragmented (%) | 1.0 |
| Missing (%) | 0.3 |
| Whole genome LAI | 24.44 |

a. BUSCO with the genome mode using the Embryopyte\_odb10 (n = 1,614) dataset.

### 3.2 Nuclear genome annotation

We identified repetitive sequences in the *G. hybrida* genome assembly, summarized in Table 2. In total, repetitive sequences occupied 89.0% of the *G. hybrida* genome assembly length. The most predominant repeat family in the *G. hybrida* genome was the LTR-retrotransposons (58.9%), including Copia-type of LTR-retrotransposons (41.1%) and Gypsy-type of LTR-retrotransposons (15.6%), followed by the DNA transposons (6.3%).

**Table 2.** Repeat elements in the *G. hybrida* genome assembly.

| Class | Number of elements | Total length (bp) | Percentage to the length of assembly (%) |
| --- | --- | --- | --- |
| Retroelements | 1,287,982 | 1,402,714,121 | 60.54 |
| SINEs | 19,159 | 2,785,476 | 0.12 |
| LINEs | 95,146 | 34,607,821 | 1.49 |
| LTR elements | 1,173,677 | 1,365,320,824 | 58.92 |
| (LTR/Copia) | 626,750 | 953,270,969 | 41.14 |
| (LTR/Gypsy) | 400,953 | 360,880,307 | 15.57 |
| DNA transposons | 526,305 | 145,198,007 | 6.27 |
| Rolling-circles | 20,955 | 4,963,828 | 0.21 |
| Unclassified | 1,897,350 | 449,420,001 | 19.40 |
| Satellites | 15,594 | 5,082,880 | 0.22 |
| Simple repeats | 317,822 | 17,636,602 | 0.76 |
| Low complexity | 29,679 | 1,510,851 | 0.07 |
| Total base masked | - | 2,062,143,287 | 89.00 |

Repeat elements were classified by RepeatModeler. 'Unclassified' refers to the repeat elements which were failed to classify to the specific repeat class.

Protein-coding genes in the *G. hybrida* genome assembly were predicted through ab initio prediction using BRAKER3^54–56^ with the alignment of RNA-seq from *G. hybrida* and the alignment of protein sequences from other plant species, and through homology-based prediction using GeMoMa^59, 60^ with protein sequences from three Asteraceae species. Initially, a total of 75,182 genes were predicted by BRAKER3. BRAKER3 predicted a large number of genes, and these might include false positive genes. Therefore, we classified these predicted genes into HC, LC, and TE genes based on the homology and the gene expression evidence. As a result, 27,469 genes were classified as HC with BUSCO completeness of 96.6% (Table S6). The total number of LC and TE genes were 26,415 and 21,298, and BUSCO completeness in LC and TE genes were 0.2% and 1.3%, respectively (Table S6). The number of HC genes was considerably lower compared to the initial result, but BUSCO completeness of the HC genes did not show a significant change, indicating the reliability of the classification. GeMoMa predicted a total number of 74,140 genes, of which 34,169, 17,109, and 25,862 genes were classified as HC, LC, and TE, respectively (Table S6). BUSCO completeness in HC, LC, and TE genes were 96.0%, 0.2%, and 1.3%, respectively.

To improve completeness of the gene prediction, we integrated the HC genes predicted by BRAKER3 and GeMoMa. Finally, the number of protein-coding genes in the *G. hybrida* genome assembly was 36,160, with BUSCO completeness of 97.3% (Table 3). Of these, 34,375 genes were assigned functional annotations based on a search against databases.

**Table 3.** Statistics of the protein-coding gene predictions of the *G.hybrida* genome assembly.

|  |  |
| --- | --- |
| Number of sequences | 36,160 |
| Total length of CDS<sup>a</sup> (bases) | 40,681,517 |
| Average length of CDS (bases) | 1,125 |
| Maximum length of CDS (bases) | 16,329 |
| Minimum length of CDS (bases) | 63 |
| N50 of CDS (bases) | 1,500 |
| A (bases) | 11,847,296 |
| T (bases) | 11,066,910 |
| G (bases) | 9,573,233 |
| C (bases) | 8,194,078 |
| GC content (%) | 43.7 |
| BUSCO<sup>b</sup> |  |
| Complete (%) | 97.3 |
| (Complete and single-copy [%]) | 90.1 |
| (Complete and duplicated [%]) | 7.2 |
| Fragmented (%) | 0.9 |
| Missing (%) | 1.8 |

a. Coding sequence.
b. BUSCO with the protein mode using the Embryopete\_odb10 (n = 1,614) dataset.

In addition, 11,572 ncRNA genes including tRNA, rRNA, miRNA, snRNA, and sno RNA were predicted (Table 4). There were 607 tRNAs, 8,328 rRNAs including 5S rRNAs, large subunit rRNAs, and small subunit rRNAs, 2,009 snoRNAs, and 291 snRNAs.

**Table 4.** The number of predicted non-coding RNA genes in the *G. hybrida* genome assembly.

| RNA type | The number of sequences |
| --- | --- |
| tRNA | 607 |
| rRNA | 8,328 |
| miRNA | 193 |
| snoRNA | 2,009 |
| snRNA | 291 |
| sRNA | 1 |
| Ribozyme | 7 |
| Antisense | 17 |
| Other genes | 19 |
| Intron | 100 |
| Total | 11,572 |

Predicted genes were classified into RNA types according to reference sequences in Rfam.

### 3.3 Organelle genome assembly and annotation

The chloroplast and the mitochondrial genomes of *G. hybrida* were assembled using HiFi reads. For the chloroplast genome, single-circular contigs with 151,898 bp of length and 37.7% of GC content was generated (Table 5). This sequence had one large single-copy region, one small single-copy region, and two inverted repeat regions (Figure 3A). As a result of gene annotation based on the reference gene alignment and de novo analysis, 85 protein-coding, 37 tRNA, and 8 rRNA genes were annotated in the chloroplast genome sequence (Table 5, Table S7, and Figure 3A). For the mitochondrial genome, single-circular contigs with a length of 363,511 bp and 45.2% of GC content was generated (Table 5). In the mitochondrial genome sequence, 36 protein-coding, 21 tRNA, and 3 rRNA genes were annotated (Table 5, Table S8, and Figure 3B).

**Figure 3.**
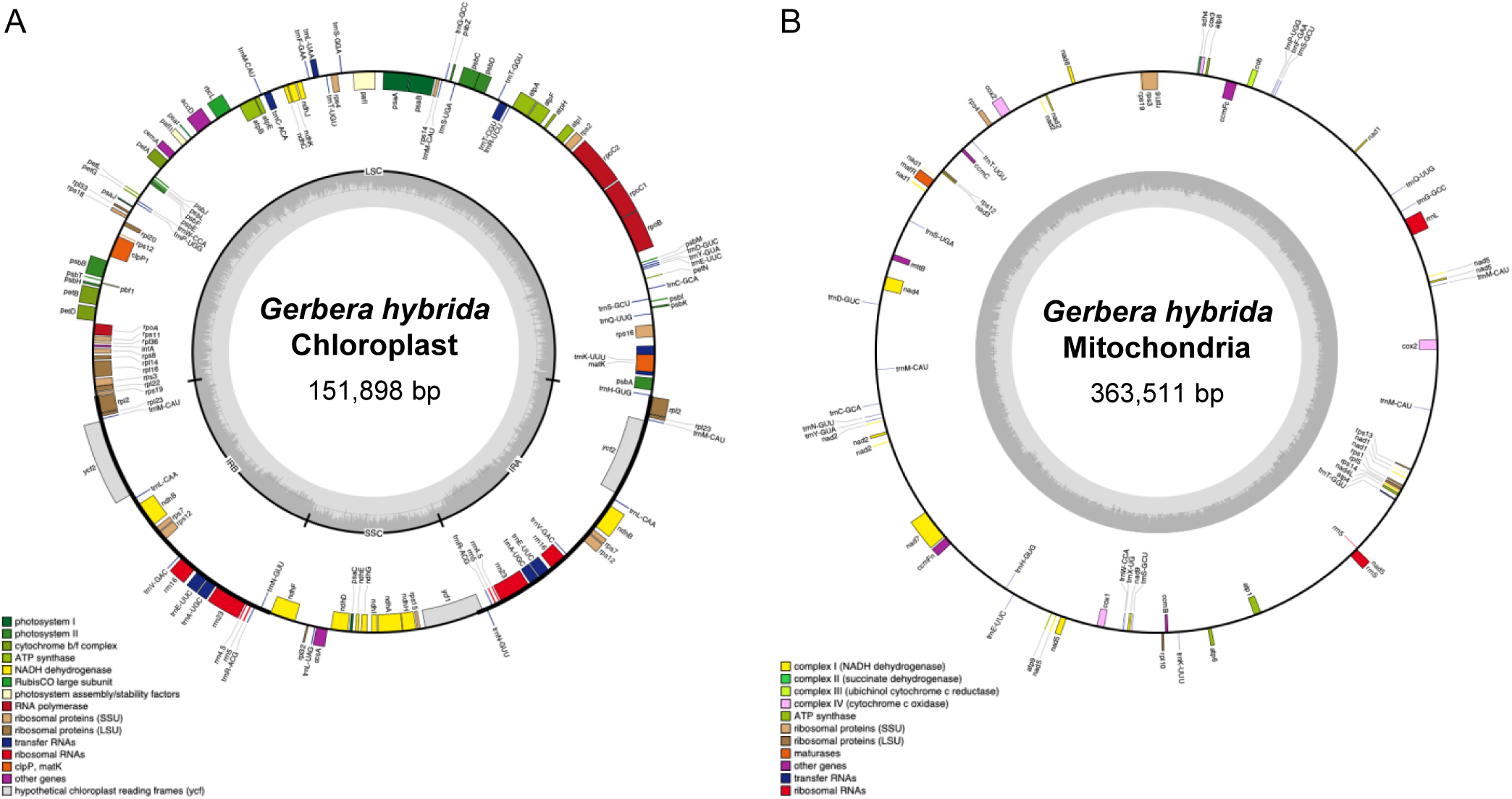
The chloroplast and the mitochondrial genomes of *G. hybrida*. The assemblies and the gene annotations of the chloroplast (A) and the mitochondrial genomes (B). Each rectangle in the outer circle indicates a gene and is colored according to the gene group. The gray bar plot in the inner circle represents the GC content. The chloroplast genome (A) contains large-single copy (LSC), small-single copy (SSC), and two inverted repeats (IRA and IRB) regions shown in the inner circle.

**Table 5.** Assemblies and gene annotations for the chloroplast and the mitochondrial genomes of *G. hybrida*.

|  | Chloroplast | Mitochondria |
| --- | --- | --- |
| **Assembly** |  |  |
| Number of sequences | 1 | 1 |
| Length (bases) | 151,898 | 363,511 |
| GC content (%) | 37.7 | 45.2 |
| **Gene** |  |  |
| Protein-coding | 85 | 36 |
| tRNA | 37 | 21 |
| rRNA | 8 | 3 |
| Total | 130 | 60 |

The complete chloroplast genome of the genus *Gerbera* has already been reported for two species.^87, 88^ Among these, the chloroplast genome of *G. hybrida* is completely identical in nucleotide sequence to that of one of its parents, *G. jamesonii*. On the other hand, at least one SNP was found between *G. jamesonii* and *G. viridifolia*.^89^ The initial hybridization of *G. hybrida* was a bidirectional cross between *G. viridifolia* and *G. jamesonii*.^4^ It remains unclear which direction of the cross gave rise to the current cultivars. The fact that *G. viridifolia* was lost shortly after the initial hybridization suggests the possibility that *G. jamesonii* became the seed parent through subsequent backcrossing. In any case, *G. jamesonii* likely contributed as the seed parent to present-day cultivars. Furthermore, investigating polymorphisms among multiple individuals within *G. hybrida* will be necessary for a more detailed understanding of the plant’s origin and development.

### 3.4 Phylogenetic analysis and genome synteny

A total of 861 orthologous groups were constructed using OrthoPhy. Phylogenetic analysis of the Asteraceae revealed that *G.hybrida* diverges at the basal position of the family (Figure 4A). This result supports previous findings based on morphological characteristics and chloroplast DNA, which suggest that the subfamily Mutisioideae and *Gerbera* represent basal lineages within the Asteraceae.^14, 90^

**Figure 4.**
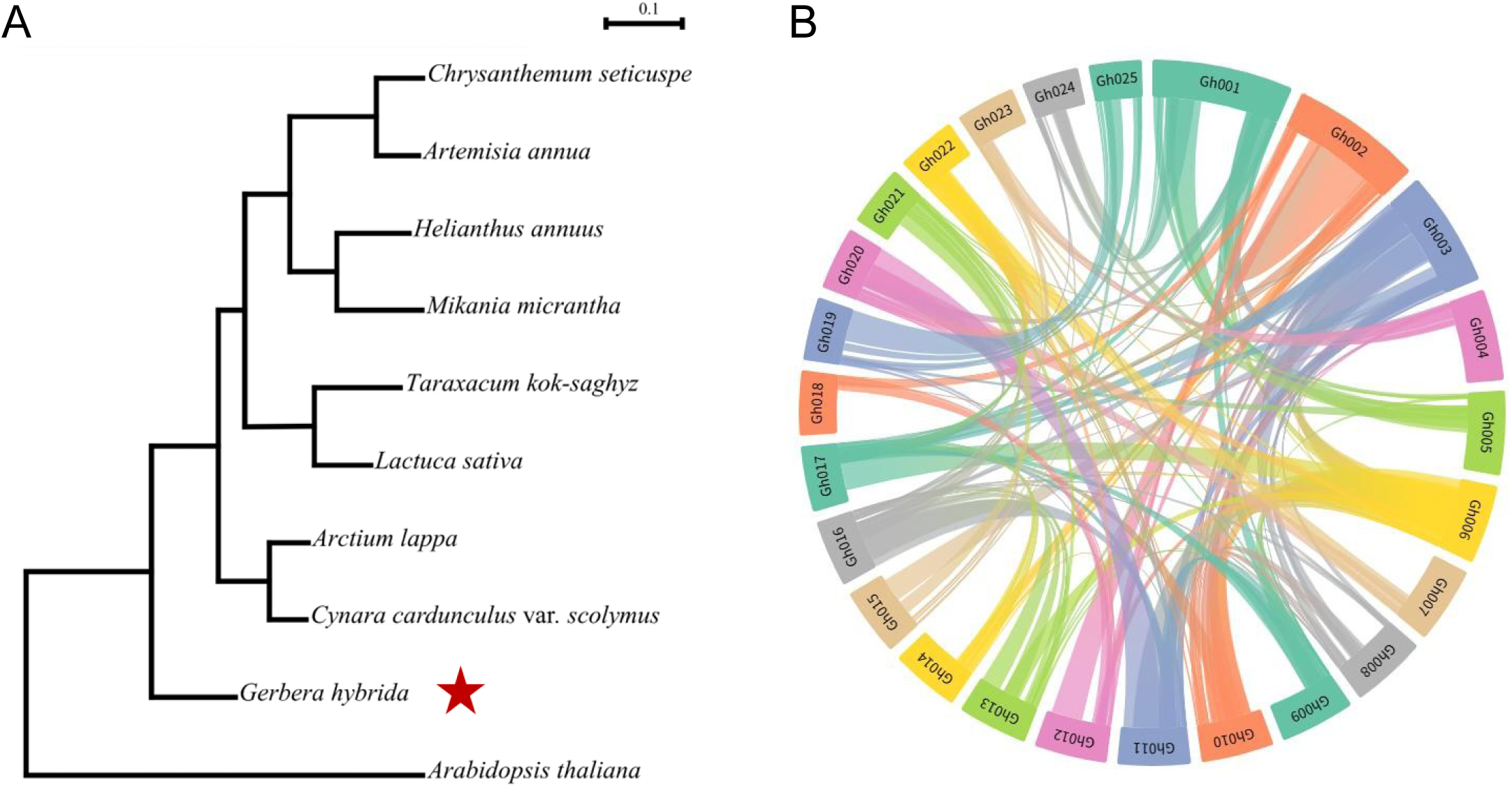
Phylogenetic relationships and genome-wide synteny figure of *G. hybrida*. (A) Concatenated tree of nine species in the Asteraceae family constructed from an orthologous protein sequences. *A. thaliana* was used as an outgroup. The scale bar means the expected number of substitution per amino acid site. *G. hybrida* is highlighted with red star. (B) Intraspecific circle synteny figure. Each collinear fragment is shown in a randomly assigned color. Gh0xx denotes the scaffold number in the *G. hybrida* genome assembly.

It has been shown that the Asteraceae experienced a shared whole-genome triplication event at its base. In fact, genome triplication has been observed in many species of the family, such as lettuce, chrysanthemum, and chicory.^11, 13, 79, 91^ Previous studies have also indicated that *G. hybrida* underwent one whole-genome triplication and one whole-genome duplication.^92^ Synteny analysis using whole-genome data revealed the complex genome structure of *G. hybrida* (Figure 4B), with some syntenic trios identified (Figure S3). This genomic complexity supports the notion that *G. hybrida* has experienced multiple genome duplication events, including the common triplication event shared among Asteraceae.

## 4. Conclusions

In this study, we report a chromosome-level genome assembly of *G. hybrida*. This represents the first reference genome for the subfamily Mutisioideae and is expected to serve as a valuable resource for elucidating the evolutionary history of the highly diverse Asteraceae family. Indeed, *G. hybrida* has been shown to have diverged earlier than other species in the Asteraceae. In cultivated gerbera, which has been bred through interspecific hybridization, this high-quality genome will facilitate the identification of gerbera-specific genes and their functions. It will also strongly support molecular breeding efforts, including genetic modification and genome editing. Overall, this genomic resource provides a fundamental genetic platform not only for gerbera but also for advancing research across the Asteraceae family.

## Acknowledgements

We would like to thank Takaharu Kimura and Akiko Watanabe of Kazusa DNA Research Institute for technical assistance. This work was supported by JSPS KAKENHI Grant Numbers JP22H04925 (PAGS), P22K05628, 22H05172, and 22H05181 and Kazusa DNA Research Institute Foundation.

## Conflict of interest

The authors declare that there is no conflict of interest.

## Author contribution

R.S., T.N., and A.Tm.: Conceptualization and study design. R.S., A.Ty., H.T., T.F. and A.Tm.: Sample preparation and experiments. Y.B.A., R.S., H.H., S.I., N.T., K.S., T.H., H.F. and M.B.: Data analysis.

Y.B.A. and R.S.: Visualization. Y.B.A. and R.S.: Writing – original draft. Y.B.A., R.S., H.H. and A.Tm.: Writing – review & editing. All authors have read and approved the manuscript.

## Data availability

Raw sequencing data used in this study have been deposited in the DNA Data Bank of Japan (DDBJ) Sequence Read Archive (DRA) under the BioProject PRJDB14615 for PacBio HiFi and Omni-C, and PRJDB35793 for RNA-seq. The accession number of each sequencing data was listed in Supplementary Table S3. The gene annotation is also available from Kazusa Genome Atlas and Plant GARDEN.

## Notes

### Competing Interest Statement

The authors have declared no competing interest.

## References

1. Yuan, H., Zhou, Q., Wani, M.A., et al. 2024. Construction of a genome-wide SSR marker library in *Gerbera hybrida*: Insights into genetic variation and germplasm resources. Sci. Hortic., 324. 112543. doi:10.1016/j.scienta.2023.112543.

2. MAFF. THE 98th STATISTICAL YEARBOOK OF MINISTRY OF AGRICULTURE, FORESTRY AND FISHERIES. https://www.maff.go.jp/e/data/stat/98th/index.html

3. Hosoguchi, T., Uchiyama, Y., Komazawa, H., Yahata, M., Shimokawa, T., and Tominaga, A. 2021. Effect of Three Types of Ion Beam Irradiation on Gerbera (*Gerbera hybrida*) In Vitro Shoots with Mutagenesis Efficiency. Plants, 10. 1480. doi:10.3390/plants10071480.

4. Hansen, H.V. 1999. A story of the cultivated Gerbera. New Plantsman, 6. 85–95.

5. Mandel, J.R., Dikow, R.B., Siniscalchi, C.M., Thapa, R., Watson, L.E., and Funk, V.A. 2019. A fully resolved backbone phylogeny reveals numerous dispersals and explosive diversifications throughout the history of Asteraceae. Proc. Natl. Acad. Sci. U.S.A., 116. 14083–14088. doi:10.1073/pnas.1903871116.

6. Panero, J.L. and Crozier, B.S. 2016. Macroevolutionary dynamics in the early diversification of Asteraceae. Mol. Phylogenet. Evol., 99. 116–132. doi:10.1016/j.ympev.2016.03.007.

7. Vijverberg, K., Welten, M., Kraaij, M., Van Heuven, B.J., Smets, E., and Gravendeel, B. 2021. Sepal Identity of the Pappus and Floral Organ Development in the Common Dandelion (*Taraxacum officinale*; Asteraceae). Plants, 10. 1682. doi:10.3390/plants10081682.

8. Xiong, W., Risse, J., Berke, L., et al. 2023. Phylogenomic analysis provides insights into MADS-box and TCP gene diversification and floral development of the Asteraceae, supported by de novo genome and transcriptome sequences from dandelion (*Taraxacum officinale*). Front. Plant Sci., 14. 1198909. doi:10.3389/fpls.2023.1198909.

9. Song, A., Su, J., Wang, H., et al. 2023. Analyses of a chromosome-scale genome assembly reveal the origin and evolution of cultivated chrysanthemum. Nature Communications, 14. doi:10.1038/s41467-023-37730-3.

10. Badouin, H., Gouzy, J., Grassa, C.J., et al. 2017. The sunflower genome provides insights into oil metabolism, flowering and Asterid evolution. Nature, 546. 148–152. doi:10.1038/nature22380.

11. Reyes-Chin-Wo, S., Wang, Z., Yang, X., et al. 2017. Genome assembly with in vitro proximity ligation data and whole-genome triplication in lettuce. Nat. Commun., 8. 14953. doi:10.1038/ncomms14953.

12. Liu, B., Yan, J., Li, W., et al. 2020. *Mikania micrantha* genome provides insights into the molecular mechanism of rapid growth. Nat. Commun., 11. 340. doi:10.1038/s41467-019-13926-4.

13. Nakano, M., Hirakawa, H., Fukai, E., et al. 2021. A chromosome-level genome sequence of *Chrysanthemum seticuspe*, a model species for hexaploid cultivated chrysanthemum. *Commun*. Biol., 4. 1167. doi:10.1038/s42003-021-02704-y.

14. Panero, J.L., Freire, S.E., Ariza Espinar, L., Crozier, B.S., Barboza, G.E., and Cantero, J.J. 2014. Resolution of deep nodes yields an improved backbone phylogeny and a new basal lineage to study early evolution of Asteraceae. Mol. Phylogenet. Evol., 80. 43–53. doi:10.1016/j.ympev.2014.07.012.

15. Kim, H.-G., Loockerman, D.J., and Jansen, R.K. 2002. Systematic Implications of ndhF Sequence Variation in the Mutisieae (Asteraceae). Syst. Bot. 598–609.

16. Teeri, T.H., Elomaa, P., Kotilainen, M., and Albert, V.A. 2006. Mining plant diversity: Gerbera as a model system for plant developmental and biosynthetic research. BioEssays, 28. 756–767. doi:10.1002/bies.20439.

17. Laitinen, R.A., Broholm, S., Albert, V.A., Teeri, T.H., and Elomaa, P. 2006. Patterns of MADS-box gene expression mark flower-type development in *Gerbera hybrida*(Asteraceae). BMC Plant Biol., 6. 11. doi:10.1186/1471-2229-6-11.

18. Lin, X., Huang, S., Huang, G., Chen, Y., Wang, X., and Wang, Y. 2021. 14-3-3 Proteins Are Involved in BR-Induced Ray Petal Elongation in *Gerbera hybrida*. Front. Plant Sci., 12. 718091. doi:10.3389/fpls.2021.718091.

19. Ren, G., Li, L., Patra, B., et al. 2023. The transcription factor GhTCP7 suppresses petal expansion by interacting with the WIP-type zinc finger protein GhWIP2 in *Gerbera hybrida*. J. Exp. Bot., 74. 4093– 4109. doi:10.1093/jxb/erad152.

20. Laitinen, R.A.E., Ainasoja, M., Broholm, S.K., Teeri, T.H., and Elomaa, P. 2008. Identification of target genes for a MYB-type anthocyanin regulator in *Gerbera hybrida*. J. Exp. Bot., 59. 3691–3703. doi:10.1093/jxb/ern216.

21. Zhong, C., Tang, Y., Pang, B., et al. 2020. The R2R3-MYB transcription factor GhMYB1a regulates flavonol and anthocyanin accumulation in *Gerbera hybrida*. Hortic. Res., 7. doi:10.1038/s41438-020-0296-2.

22. Zhu, L., Zhang, T., and Teeri, T.H. 2021. Tetraketide α-pyrone reductases in sporopollenin synthesis pathway in *Gerbera hybrida*: diversification of the minor function. Hortic. Res., 8. doi:10.1038/s41438-021-00642-8.

23. Zhu, L., Pietiäinen, M., Kontturi, J., Turkkelin, A., Elomaa, P., and Teeri, T.H. 2022. Polyketide reductases in defense-related parasorboside biosynthesis in *Gerbera hybrida* share processing strategies with microbial polyketide synthase systems. New Phytol., 236. 296–308. doi:10.1111/nph.18328.

24. Bhattarai, K., Sharma, S., Verma, S., et al. 2023. Construction of a genome-wide genetic linkage map and identification of quantitative trait loci for powdery mildew resistance in *Gerbera* daisy. Front. Plant Sci., 13. 1072717. doi:10.3389/fpls.2022.1072717.

25. Luo, Q., Huang, G., Lin, X., Wang, X., and Wang, Y. 2025. Genome-wide identification, characterization, and expression analysis of BZR transcription factor family in *Gerbera hybrida*. BMC Plant Biol., 25. 143. doi:10.1186/s12870-025-06177-7.

26. Baid G, Cook DE, Shafin K, et al. DeepConsensus improves the accuracy of sequences with a gap-aware sequence transformer. Nat Biotechnol. 2023;41:232–238. doi:10.1038/s41587-022-01435-7

27. Sim, S.B., Corpuz, R.L., Simmonds, T.J., and Geib, S.M. 2022. HiFiAdapterFilt, a memory efficient read processing pipeline, prevents occurrence of adapter sequence in PacBio HiFi reads and their negative impacts on genome assembly. BMC Genomics, 23. 157. doi:10.1186/s12864-022-08375-1.

28. Martin, M. 2011. Cutadapt removes adapter sequences from high-throughput sequencing reads. EMBnet.journal, 17. 10–12. doi:10.14806/EJ.17.1.200.

29. Chen, S., Zhou, Y., Chen, Y., and Gu, J. 2018. fastp: an ultra-fast all-in-one FASTQ preprocessor. Bioinformatics, 34. i884–i890. doi:10.1093/bioinformatics/bty560.

30. Rhie, A., Walenz, B.P., Koren, S., and Phillippy, A.M. 2020. Merqury: reference-free quality, completeness, and phasing assessment for genome assemblies. Genome Biol., 21. 245. doi:10.1186/s13059-020-02134-9.

31. Ranallo-Benavidez, T.R., Jaron, K.S., and Schatz, M.C. 2020. GenomeScope 2.0 and Smudgeplot for reference-free profiling of polyploid genomes. Nat. Commun., 11. 1432. doi:10.1038/s41467-020-14998-3.

32. Cheng, H., Concepcion, G.T., Feng, X., Zhang, H., and Li, H. 2021. Haplotype-resolved de novo assembly using phased assembly graphs with hifiasm. Nat. Methods, 18. 170–175. doi:10.1038/s41592-020-01056-5.

33. Cheng, H., Jarvis, E.D., Fedrigo, O., et al. 2022. Haplotype-resolved assembly of diploid genomes without parental data. Nat. Biotechnol., 40. 1332–1335. doi:10.1038/s41587-022-01261-x.

34. Li, H. 2018. Minimap2: pairwise alignment for nucleotide sequences. Bioinformatics, 34. 3094–3100. doi:10.1093/bioinformatics/bty191.

35. Li, H. 2021. New strategies to improve minimap2 alignment accuracy. Bioinformatics, 37. 4572–4574. doi:10.1093/bioinformatics/btab705.

36. Roach, M.J., Schmidt, S.A., and Borneman, A.R. 2018. Purge Haplotigs: allelic contig reassignment for third-gen diploid genome assemblies. BMC Bioinformatics, 19. 460. doi:10.1186/s12859-018-2485-7.

37. Li, H. 2013. Aligning sequence reads, clone sequences and assembly contigs with BWA-MEM. arXiv [q-bio.GN].

38. Open2C, Abdennur, N., Fudenberg, G., et al. 2024. Pairtools: From sequencing data to chromosome contacts. PLoS Comput. Biol., 20. e1012164. doi:10.1371/journal.pcbi.1012164.

39. Ghurye, J., Pop, M., Koren, S., Bickhart, D., and Chin, C.-S. 2017. Scaffolding of long read assemblies using long range contact information. BMC Genomics, 18. 527. doi:10.1186/s12864-017-3879-z.

40. Ghurye, J., Rhie, A., Walenz, B.P., et al. 2019. Integrating Hi-C links with assembly graphs for chromosome-scale assembly. PLoS Comput. Biol., 15. e1007273. doi:10.1371/journal.pcbi.1007273.

41. Quinlan, A.R. and Hall, I.M. 2010. BEDTools: a flexible suite of utilities for comparing genomic features. Bioinformatics, 26. 841–842. doi:10.1093/bioinformatics/btq033.

42. Durand, N.C., Shamim, M.S., Machol, I., et al. 2016. Juicer provides a one-click system for analyzing loop-resolution hi-C experiments. Cell Syst., 3. 95–98. doi:10.1016/j.cels.2016.07.002.

43. Ramírez, F., Ryan, D.P., Grüning, B., et al. 2016. deepTools2: a next generation web server for deep-sequencing data analysis. Nucleic Acids Res., 44. W160–5. doi:10.1093/nar/gkw257.

44. Cabanettes, F. and Klopp, C. 2018. D-GENIES: dot plot large genomes in an interactive, efficient and simple way. PeerJ, 6. e4958. doi:10.7717/peerj.4958.

45. Brown, M.R., Gonzalez de La Rosa, P., and Blaxter, M. 2025. Tidk: A toolkit to rapidly identify telomeric repeats from genomic datasets. Bioinformatics. doi:10.1093/bioinformatics/btaf049.

46. Manni, M., Berkeley, M.R., Seppey, M., Simão, F.A., and Zdobnov, E.M. 2021. BUSCO Update: Novel and Streamlined Workflows along with Broader and Deeper Phylogenetic Coverage for Scoring of Eukaryotic, Prokaryotic, and Viral Genomes. Mol. Biol. Evol., 38. 4647–4654. doi:10.1093/molbev/msab199.

47. Ou, S., Chen, J., and Jiang, N. 2018. Assessing genome assembly quality using the LTR Assembly Index (LAI). Nucleic Acids Res., 46. e126. doi:10.1093/nar/gky730.

48. Ou, S. and Jiang, N. 2018. LTR_retriever: A highly accurate and sensitive program for identification of long terminal repeat retrotransposons. Plant Physiol., 176. 1410–1422. doi:10.1104/pp.17.01310.

49. Flynn, J.M., Hubley, R., Goubert, C., et al. 2020. RepeatModeler2 for automated genomic discovery of transposable element families. Proc. Natl. Acad. Sci. U.S.A., 117. 9451–9457. doi:10.1073/pnas.1921046117.

50. Bao, W., Kojima, K.K., and Kohany, O. 2015. Repbase Update, a database of repetitive elements in eukaryotic genomes. Mob. DNA, 6. 11. doi:10.1186/s13100-015-0041-9.

51. Storer, J., Hubley, R., Rosen, J., Wheeler, T.J., and Smit, A.F. 2021. The Dfam community resource of transposable element families, sequence models, and genome annotations. Mob. DNA, 12. 2. doi:10.1186/s13100-020-00230-y.

52. UniProt Consortium. 2023. UniProt: The universal protein knowledgebase in 2023. Nucleic Acids Res., 51. D523–D531. doi:10.1093/nar/gkac1052.

53. Buchfink, B., Reuter, K., and Drost, H.-G. 2021. Sensitive protein alignments at tree-of-life scale using DIAMOND. Nat. Methods, 18. 366–368. doi:10.1038/s41592-021-01101-x.

54. Gabriel, L., Brůna, T., Hoff, K.J., et al. 2024. BRAKER3: Fully automated genome annotation using RNA-seq and protein evidence with GeneMark-ETP, AUGUSTUS, and TSEBRA. Genome Res., 34. 769–777. doi:10.1101/gr.278090.123.

55. Stanke, M., Schöffmann, O., Morgenstern, B., and Waack, S. 2006. Gene prediction in eukaryotes with a generalized hidden Markov model that uses hints from external sources. BMC Bioinformatics, 7. 62. doi:10.1186/1471-2105-7-62.

56. Stanke, M., Diekhans, M., Baertsch, R., and Haussler, D. 2008. Using native and syntenically mapped cDNA alignments to improve de novo gene finding. Bioinformatics, 24. 637–644. doi:10.1093/bioinformatics/btn013.

57. Dobin, A., Davis, C.A., Schlesinger, F., et al. 2013. STAR: ultrafast universal RNA-seq aligner. Bioinformatics, 29. 15–21. doi:10.1093/bioinformatics/bts635.

58. Tegenfeldt, F., Kuznetsov, D., Manni, M., Berkeley, M., Zdobnov, E.M., and Kriventseva, E.V. 2025. OrthoDB and BUSCO update: annotation of orthologs with wider sampling of genomes. Nucleic Acids Res., 53. D516–D522. doi:10.1093/nar/gkae987.

59. Keilwagen, J., Wenk, M., Erickson, J.L., Schattat, M.H., Grau, J., and Hartung, F. 2016. Using intron position conservation for homology-based gene prediction. Nucleic Acids Res., 44. e89. doi:10.1093/nar/gkw092.

60. Keilwagen, J., Hartung, F., Paulini, M., Twardziok, S.O., and Grau, J. 2018. Combining RNA-seq data and homology-based gene prediction for plants, animals and fungi. BMC Bioinformatics, 19. 189. doi:10.1186/s12859-018-2203-5.

61. UniProt Consortium. 2025. UniProt: The universal protein knowledgebase in 2025. Nucleic Acids Res., 53. D609–D617. doi:10.1093/nar/gkae1010.

62. Huerta-Cepas, J., Szklarczyk, D., Heller, D., et al. 2019. eggNOG 5.0: a hierarchical, functionally and phylogenetically annotated orthology resource based on 5090 organisms and 2502 viruses. Nucleic Acids Res., 47. D309–D314. doi:10.1093/nar/gky1085.

63. Cantalapiedra, C.P., Hernández-Plaza, A., Letunic, I., Bork, P., and Huerta-Cepas, J. 2021. EggNOG-mapper v2: Functional annotation, orthology assignments, and domain prediction at the metagenomic scale. Mol. Biol. Evol., 38. 5825–5829. doi:10.1093/molbev/msab293.

64. Paysan-Lafosse, T., Andreeva, A., Blum, M., et al. 2025. The Pfam protein families database: embracing AI/ML. Nucleic Acids Res., 53. D523–D534. doi:10.1093/nar/gkae997.

65. Eddy, S.R. 2011. Accelerated profile HMM searches. PLoS Comput. Biol., 7. e1002195. doi:10.1371/journal.pcbi.1002195.

66. Patro, R., Duggal, G., Love, M.I., Irizarry, R.A., and Kingsford, C. 2017. Salmon provides fast and bias-aware quantification of transcript expression. Nat. Methods, 14. 417–419. doi:10.1038/nmeth.4197.

67. Pertea, G. and Pertea, M. 2020. GFF utilities: GffRead and GffCompare. F1000Res., 9. 304. doi:10.12688/f1000research.23297.2.

68. Chan, P.P., Lin, B.Y., Mak, A.J., and Lowe, T.M. 2021. tRNAscan-SE 2.0: improved detection and functional classification of transfer RNA genes. Nucleic Acids Res., 49. 9077–9096. doi:10.1093/nar/gkab688.

69. Nawrocki, E.P. and Eddy, S.R. 2013. Infernal 1.1: 100-fold faster RNA homology searches. Bioinformatics, 29. 2933. doi:10.1093/bioinformatics/btt509.

70. Ontiveros-Palacios, N., Cooke, E., Nawrocki, E.P., et al. 2025. Rfam 15: RNA families database in 2025. Nucleic Acids Res., 53. D258–D267. doi:10.1093/nar/gkae1023.

71. Krzywinski, M., Schein, J., Birol, İ., et al. 2009. Circos: An information aesthetic for comparative genomics. Genome Research, 19. 1639–1645. doi:10.1101/gr.092759.109.

72. Zhou, C., Brown, M., Blaxter, M., The Darwin Tree of Life Project Consortium, McCarthy, S.A., and Durbin, R. 2024. Oatk: a de novo assembly tool for complex plant organelle genomes. bioRxiv. doi:10.1101/2024.10.23.619857.

73. Tillich, M., Lehwark, P., Pellizzer, T., et al. 2017. GeSeq - versatile and accurate annotation of organelle genomes. Nucleic Acids Res., 45. W6–W11. doi:10.1093/nar/gkx391.

74. Kent, W.J. 2002. BLAT--the BLAST-like alignment tool. Genome Res., 12. 656–664. doi:10.1101/gr.229202.

75. Laslett, D. and Canback, B. 2004. ARAGORN, a program to detect tRNA genes and tmRNA genes in nucleotide sequences. Nucleic Acids Res., 32. 11–16. doi:10.1093/nar/gkh152.

76. Greiner, S., Lehwark, P., and Bock, R. 2019. OrganellarGenomeDRAW (OGDRAW) version 1.3.1: expanded toolkit for the graphical visualization of organellar genomes. Nucleic Acids Res., 47. W59– W64. doi:10.1093/nar/gkz238.

77. Marçais, G., Delcher, A.L., Phillippy, A.M., Coston, R., Salzberg, S.L., and Zimin, A. 2018. MUMmer4: A fast and versatile genome alignment system. PLOS Computational Biology, 14. e1005944. doi:10.1371/journal.pcbi.1005944.

78. Hirakawa, H., Sumitomo, K., Hisamatsu, T., et al. 2019. De novo whole-genome assembly in *Chrysanthemum seticuspe*, a model species of Chrysanthemums, and its application to genetic and gene discovery analysis. DNA Research, 26. 195–203. doi:10.1093/dnares/dsy048.

79. Shen, Q., Zhang, L., Liao, Z., et al. 2018. The Genome of Artemisia annua Provides Insight into the Evolution of Asteraceae Family and Artemisinin Biosynthesis. Molecular Plant, 11. 776–788. doi:10.1016/j.molp.2018.03.015.

80. Fan, W., Wang, S., Wang, H., et al. 2022. The genomes of chicory, endive, great burdock and yacon provide insights into Asteraceae palaeo-polyploidization history and plant inulin production. Mol. Ecol. Resour., 22. 3124–3140. doi:10.1111/1755-0998.13675.

81. Lamesch, P., Berardini, T.Z., Li, D., et al. 2012. The Arabidopsis Information Resource (TAIR): improved gene annotation and new tools. Nucleic Acids Research, 40. D1202–D1210. doi:10.1093/nar/gkr1090.

82. Watanabe, T., Kure, A., and Horiike, T. 2023. OrthoPhy: A Program to Construct Ortholog Data Sets Using Taxonomic Information. Genome Biol. Evol., 15. doi:10.1093/gbe/evad026.

83. Wang, Y., Tang, H., DeBarry, J.D., et al. 2012. MCScanX: a toolkit for detection and evolutionary analysis of gene synteny and collinearity. Nucleic Acids Res., 40. e49–e49. doi:10.1093/nar/gkr1293.

84. Camacho, C., Coulouris, G., Avagyan, V., et al. 2009. BLAST+: architecture and applications. BMC Bioinformatics, 10. 421. doi:10.1186/1471-2105-10-421.

85. Bandi, V., Gutwin, C., Siri, J.N., Neufeld, E., Sharpe, A., and Parkin, I. 2022. Visualization Tools for Genomic Conservation. Plant Bioinformatics: Methods and Protocols. 285–308. doi:10.1007/978-1-0716-2067-0_16.

86. Marie, D. and Brown, S.C. 1993. A cytometric exercise in plant DNA histograms, with 2C values for 70 species. Biol. Cell, 78. 41–51. doi:10.1016/0248-4900(93)90113-s.

87. Zhang, Y.-Y., Liu, F., Wang, X., et al. 2019. Characterization of the complete chloroplast genome of *Gerbera jamesonii* Bolus in China and phylogenetic relationships. Mitochondrial DNA B Resour., 4. 2706–2707. doi:10.1080/23802359.2019.1644230.

88. Sheng, W. 2025. The complete chloroplast genome of *Gerbera piloselloides* (L.) Cass., 1820 (Carduoideae, Asteraceae) and its phylogenetic analysis. Open Life Sci., 20. 20251070. doi:10.1515/biol-2025-1070.

89. Xu, X., Zheng, W., Funk, V.A., Li, K., Zhang, J., and Wen, J. 2018. Home at last III: Transferring Uechtritzia and Asian Gerbera species into Oreoseris (Compositae, Mutisieae). PhytoKeys, 96. 1–19. doi:10.3897/phytokeys.96.23142.

90. Katinas, L., Pruski, J., Sancho, G., and Tellería, M.C. 2008. The Subfamily Mutisioideae (Asteraceae). Bot. Rev., 74. 469–716. doi:10.1007/s12229-008-9016-6.

91. Yang, Y., Li, S., Xing, Y., et al. 2022. The first high-quality chromosomal genome assembly of a medicinal and edible plant *Arctium lappa*. Mol. Ecol. Resour., 22. 1493–1507. doi:10.1111/1755-0998.13547.

92. Barker, M.S., Kane, N.C., Matvienko, M., et al. 2008. Multiple Paleopolyploidizations during the Evolution of the Compositae Reveal Parallel Patterns of Duplicate Gene Retention after Millions of Years. Mol. Biol. Evol., 25. 2445–2455. doi:10.1093/molbev/msn187.

