## Supplementary_Figure for "Chromosome-level genome assembly of the gerbera (*Gerbera hybrida*) using HiFi long-read and Hi-C technologies"

### Slide 1
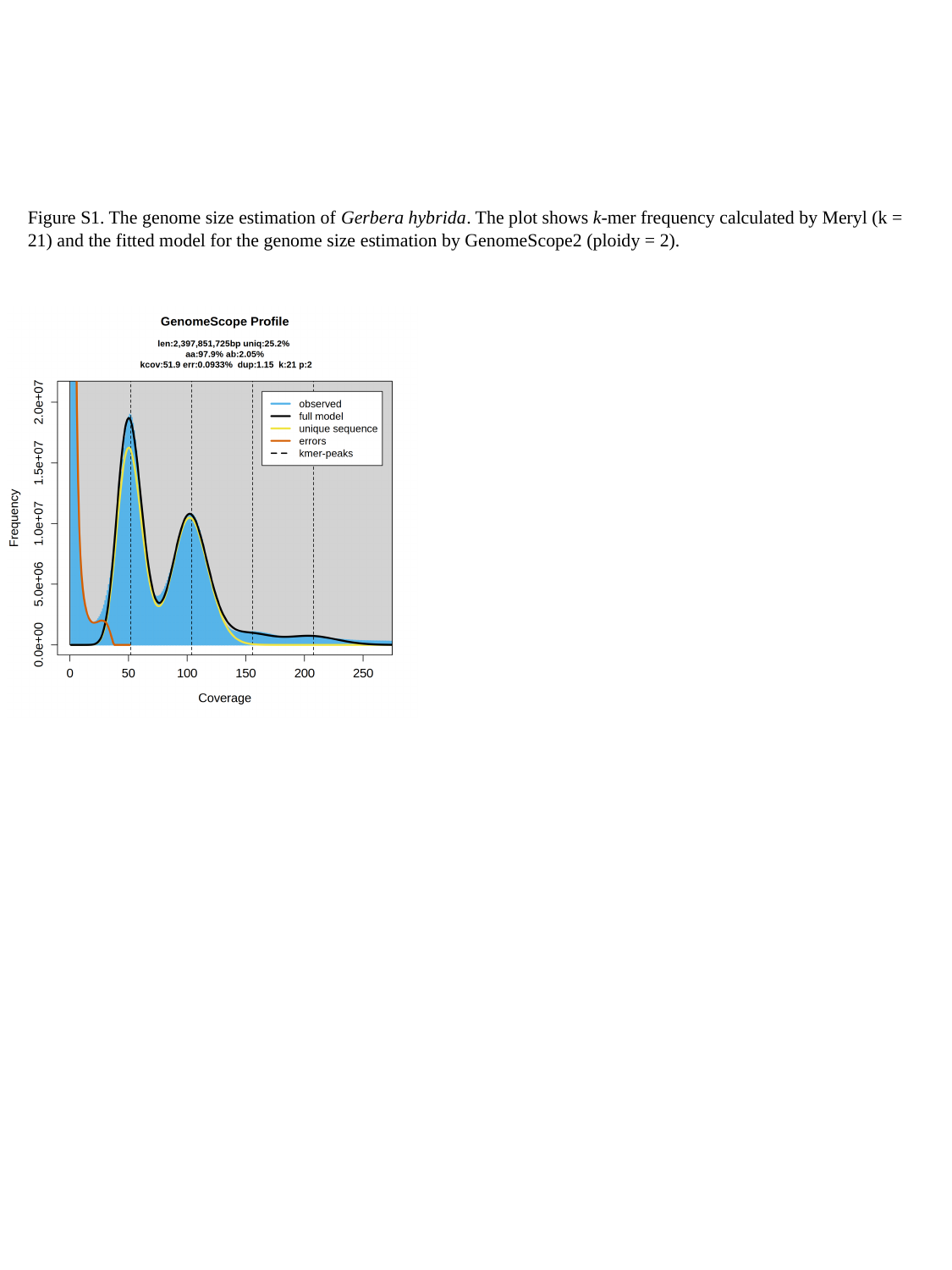

Figure S1. The genome size estimation of Gerbera hybrida. The plot shows k-mer frequency calculated by Meryl (k = 21) and the fitted model for the genome size estimation by GenomeScope2 (ploidy = 2).

### Slide 2
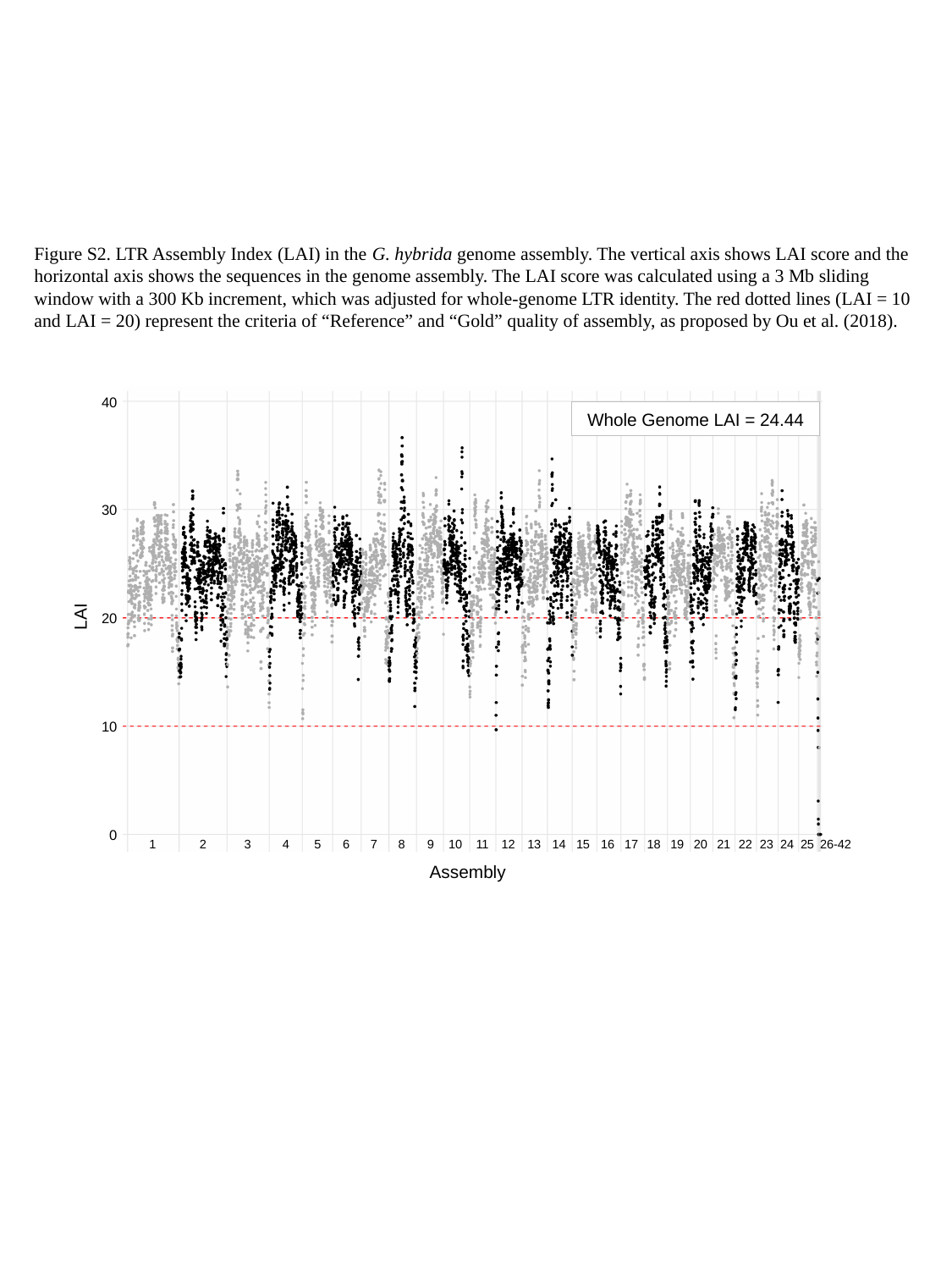

Figure S2. LTR Assembly Index (LAI) in the G. hybrida genome assembly. The vertical axis shows LAI score and the horizontal axis shows the sequences in the genome assembly. The LAI score was calculated using a 3 Mb sliding window with a 300 Kb increment, which was adjusted for whole-genome LTR identity. The red dotted lines (LAI = 10 and LAI = 20) represent the criteria of “Reference” and “Gold” quality of assembly, as proposed by Ou et al. (2018).
40
Whole Genome LAI = 24.44
30
LAI
20
10
0
1
2
3
4
5
6
7
8
9
10
11
12
13
14
15
16
17
18
19
20
21
22
23
24
25
26-42
Assembly

### Slide 3
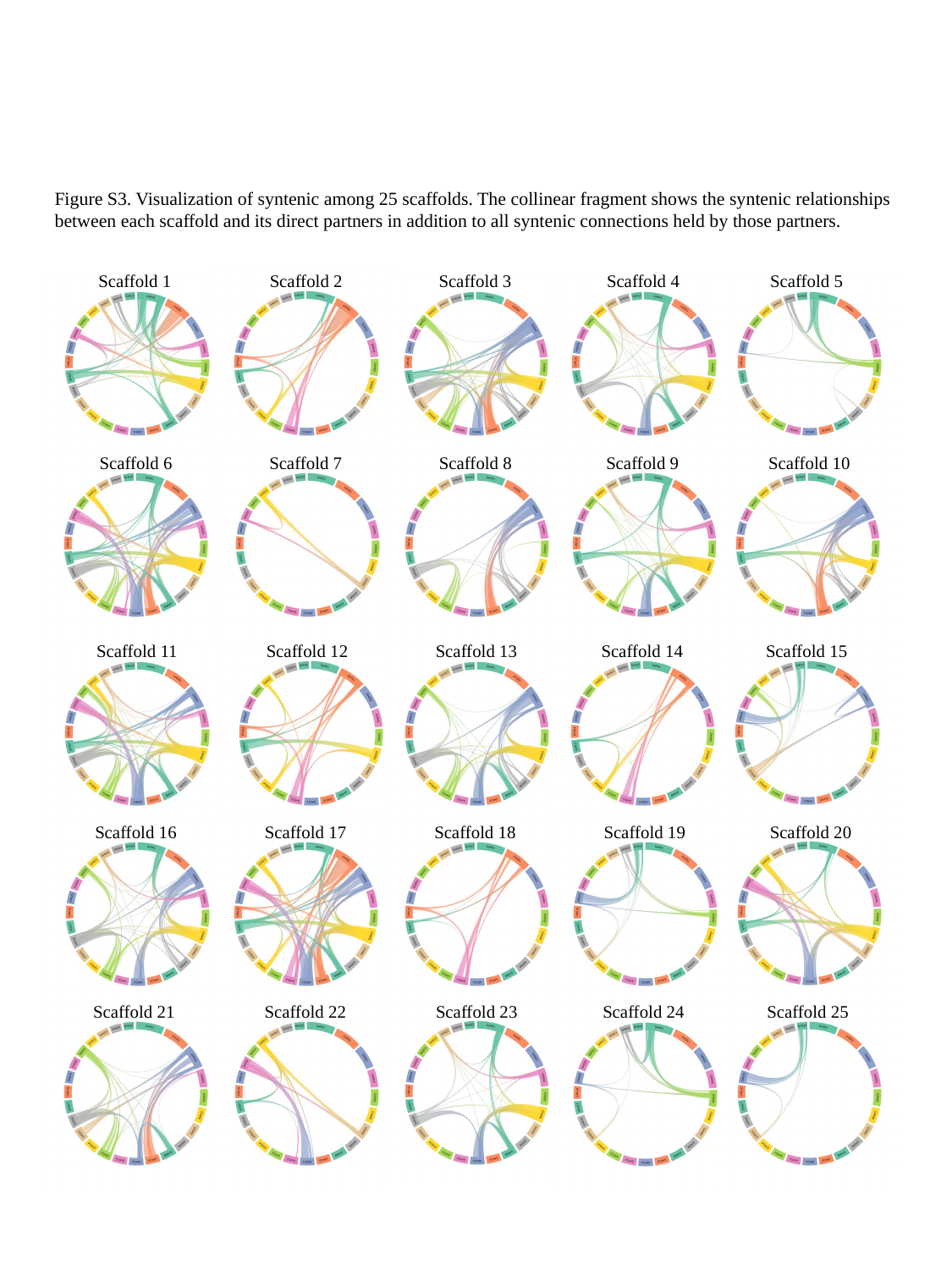

Figure S3. Visualization of syntenic among 25 scaffolds. The collinear fragment shows the syntenic relationships between each scaffold and its direct partners in addition to all syntenic connections held by those partners.
Scaffold 1
Scaffold 2
Scaffold 3
Scaffold 4
Scaffold 5
Scaffold 9
Scaffold 10
Scaffold 8
Scaffold 7
Scaffold 6
Scaffold 15
Scaffold 13
Scaffold 14
Scaffold 12
Scaffold 11
Scaffold 16
Scaffold 17
Scaffold 18
Scaffold 19
Scaffold 20
Scaffold 22
Scaffold 23
Scaffold 24
Scaffold 25
Scaffold 21
